# Verapamil and its metabolite norverapamil inhibit the *Mycobacterium tuberculosis* MmpS5L5 efflux pump to potentiate the activity bedaquiline and other antitubercular drugs

**DOI:** 10.1101/2025.02.06.636695

**Authors:** Adam J Fountain, Natalie JE Waller, Chen-Yi Cheung, William Jowsey, Michael T Chrisp, Mark Troll, Paul H Edelstein, Gregory M Cook, Matthew B McNeil, Lalita Ramakrishnan

## Abstract

Bedaquiline is the cornerstone of a new regimen for the treatment of drug-resistant tuberculosis. However, its clinical use is threatened by the emergence of bedaquiline-resistant strains of *Mycobacterium tuberculosis*. Bedaquiline targets mycobacterial ATP synthase but the predominant route to clinical bedaquiline resistance is via upregulation of the MmpS5L5 efflux pump due to mutations that inactivate the transcriptional repressor Rv0678. Here, we show that the MmpS5L5 efflux pump reduces susceptibility to bedaquiline as well as its new, more potent derivative TBAJ-876 and other antimicrobial substrates, including clofazimine and the DprE1 inhibitors PBTZ-169 and OPC-167832. Furthermore, the increased resistance of Rv0678 mutants stems entirely from increased MmpS5L5 activity. These results highlight the potential of a pharmacological MmpS5L5 inhibitor to increase drug efficacy. Verapamil, primarily used as a calcium channel inhibitor, is known to inhibit diverse efflux pumps and to potentiate bedaquiline and clofazimine activity in *M. tuberculosis*. Here, we show that verapamil potentiates the activity of multiple diverse MmpS5L5 substrates. Using biochemical approaches, we demonstrate that verapamil does not exert this effect by acting as a disruptor of the protonmotive force used to power MmpS5L5, as previously proposed, suggesting that verapamil inhibits the function of the MmpS5L5 pump. Finally, norverapamil, the major verapamil metabolite, which has greatly reduced calcium channel activity, has equal potency in reducing resistance to MmpS5L5 substrates. Our findings highlight verapamil’s potential for enhancing bedaquiline TB treatment, for preventing acquired resistance to bedaquiline and other MmpS5L5 substrates, while also providing the impetus to identify additional MmpS5L5 inhibitors.

**Significance Statement:** Bedaquiline, an antitubercular drug targeting ATP synthase, forms the backbone of highly efficacious treatment regimens for drug-resistant tuberculosis. Bedaquiline resistance is emerging as a significant problem and is most commonly caused by mutations in Rv0678 that result in upregulation of the MmpS5L5 efflux pump. Here we define the contribution of the MmpS5L5 efflux pump to drug resistance in wild-type and Rv0678 mutant strains and show that the commonly used drug verapamil can inhibit MmpS5L5 activity. This suggests that this safe and inexpensive drug may be useful in enhancing bedaquiline treatment of TB and to help prevent bedaquiline acquired resistance.

## Introduction

Drug-resistant strains of *Mycobacterium tuberculosis* are a major hurdle to combating tuberculosis (TB). Globally, 3.6% of new cases and 18% of previously treated cases of tuberculosis are multi-drug resistant (MDR)-TB (1). Until recently, treatment options for MDR-TB were limited, requiring long treatments with combinations of toxic drugs, including injectable ones, with poor treatment outcomes (2). A highly effective all-oral 6-month regimen has revolutionized MDR TB treatment (3–5). Bedaquiline, a new class of antitubercular drug that targets mycobacterial ATP synthase (6), is the cornerstone of this new regimen. Alarmingly, despite its relatively recent approval in 2012, acquired bedaquiline resistance is already widely observed (7–9). Bedaquiline resistance is associated with increased rates of treatment failure and relapse (10–14) and can occur even with optimal regimen adherence (15).

Clinical resistance to bedaquiline is rarely caused by mutations in its target ATP synthase (*atpE*). Rather, it is predominantly due to the upregulation of the MmpS5L5 efflux pump operon caused by mutations that inactivate the transcriptional repressor *Rv0678* (7,16–21). MmpL5 is a member of the Mycobacterial membrane protein Large (MmpL) transporter family (22–30). Several MmpL proteins are encoded in operons containing associated small proteins, termed mycobacterial membrane protein small (MmpS) (30–33). MmpS5 and MmpL5 are proposed to form a complex, whose physiological function is export of the iron siderophores carboxymycobactin and mycobactin (32–34), but which can also efflux bedaquiline and other structurally and functionally diverse drugs (7,20,31–33,35,36). MmpS5L5 upregulation via mutations in Rv0678 may also facilitate the transition from low-level to untreatable high-level bedaquiline resistance by allowing larger populations of bacteria to survive or tolerate the drug to allow selection of high-level resistance (37). Here, we comprehensively assess the role of MmpS5L5 in resistance to multiple antitubercular drugs, and the role and mechanism of action of the commonly used drug verapamil in overcoming MmpS5L5-mediated resistance.

## Results

### The MmpS5L5 efflux pump reduces susceptibility to bedaquiline and its derivatives

Multiple studies have found that clinical or *in vitro* derived *M. tuberculosis* strains with mutations in the MmpS5L5 transcriptional repressor Rv0678 have decreased susceptibility to bedaquiline and a diverse set of additional antitubercular drugs (20,32,35,36,38). The strength of resistance varies with Rv0678 mutant genotype, with different clinical and *in vitro* derived Rv0678 mutants having between 2- and 16-fold increased bedaquiline resistance (18,39,40). As a consistent benchmark for our study, we used our previously isolated frameshift mutant (*Rv0678* G65GfsX10) (19) to test drug MICs to reported MmpS5L5 substrates. This mutant had an 8-fold increase in the bedaquiline MIC (Fig. 1A and *SI Appendix,* Table S2). It also had increased drug MICs, 8-fold and 4-fold respectively, for the more potent bedaquiline derivatives TBAJ-867 and TBAJ-587, which are in phase 1 and 2 clinical trials (41–43) (*SI Appendix*, Table S2). This is similar to a 4-fold increase for both drugs reported previously (44,45). It also displayed decreased susceptibility to clofazimine, PBTZ-169 and OPC-167832 similar to those previously reported (32,35,36)(Fig. 1A and *SI Appendix*, Table S1). These findings confirm previous reports that Rv0678 inactivation decreases susceptibility to multiple drugs.

**Figure 1.**
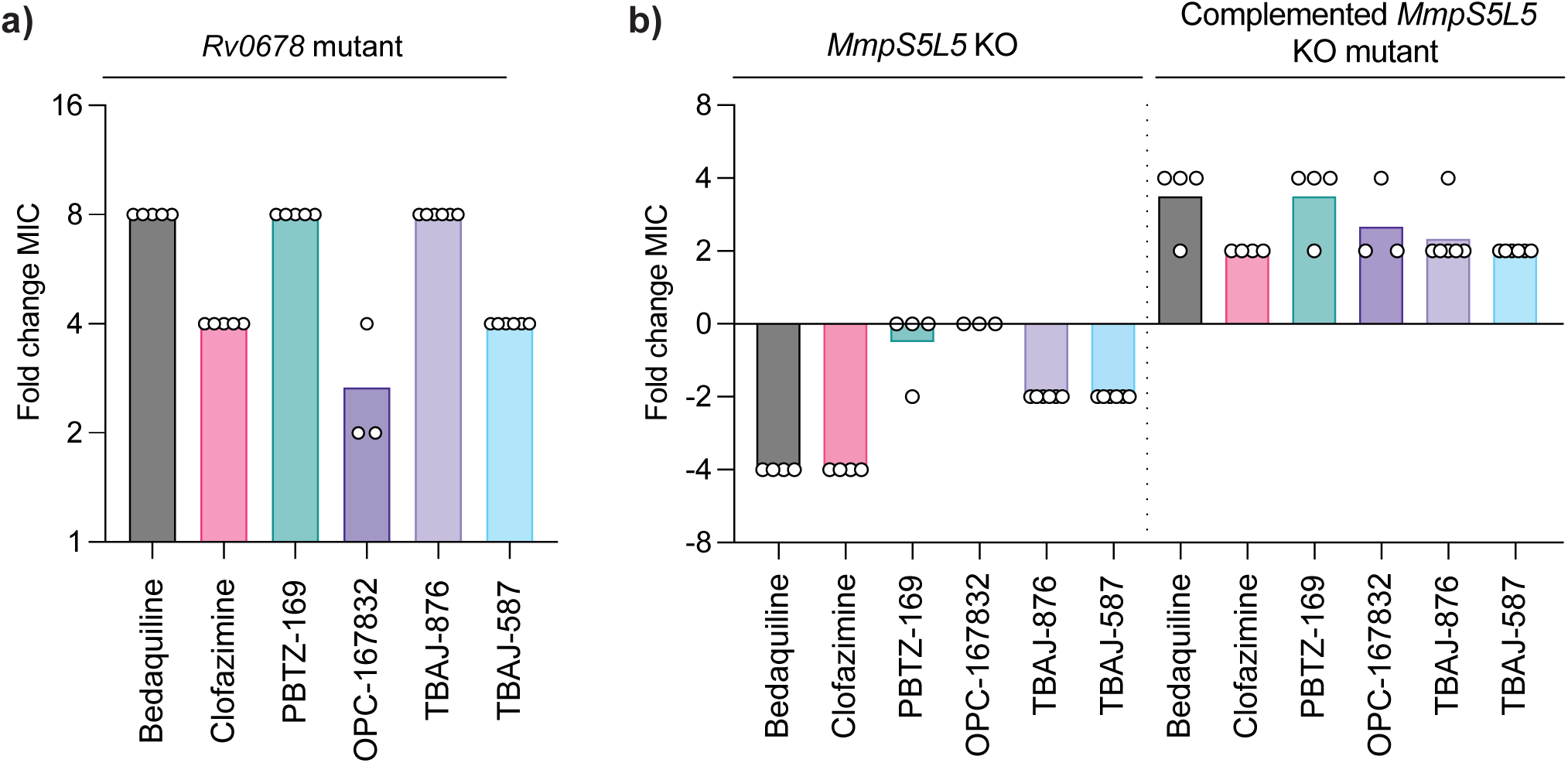
Mutations in *Rv0678* provide resistance to structurally and functionally diverse antitubercular drugs: **(a)** Fold change in MIC between wild-type *M. tuberculosis* and isogenic *Rv0678*^-^ G65GfsX10 strains against the stated drugs. Results are representative of at least three independent experiments. **(b)** Fold change in MIC between wild-type *M. tuberculosis* and isogenic Δ*S5L5*::loxP and Δ*S5L5*::loxP + pMINTF3 *M. tuberculosis MmpS5L5* strains against the stated drugs. Symbols indicate the results of technical replicates (N=3 to 6). Results are representative of at least three independent experiments. The drug MICs for the tested *M. tuberculosis* strains are shown in *SI Appendix*, Table S2.

MmpS5L5 knockout mutants have increased susceptibility to bedaquiline and clofazimine, suggesting that MmpS5L5 mediates intrinsic decreased susceptibility even in the absence of overexpression (32). We confirmed this finding and extended it to TBAJ-876 and TBAJ-587 (Fig. 1B). Complementation of the Δ*S5L5* mutant with a single-copy integrating construct of *MmpS5L5* expressed from a constitutive promoter, caused increased drug MICs (Fig. 1B). Drug MICs were 2–4 fold higher than wild-type due to higher expression levels from the non-native promoter (Fig. 1B and *SI Appendix*, Fig. S1).

In contrast, the MmpS5L5 knockout strain had no increased sensitivity to the DprE1 inhibitors OPC-167832 or PBTZ-169, but did have reduced activity against the complemented MmpS5L5 knockout strain that overexpresses MmpS5L5 (Fig. 1B). This is consistent with only modest sensitivity reported previously for PBTZ-169 (36) and suggests that while these DprE1 inhibitors are pump substrates, MmpS5L5 contributes minimally to resistance to these agents in wild-type strains.

### MmpS5L5 knockdown reverses increased resistance of Rv0678 mutant

Previous studies have linked resistance of Rv0678 mutants to MmpS5L5 upregulation (7). However, it has not been established if upregulation is the sole cause of resistance or whether Rv0678 regulates other processes contributing to resistance. To address this question, we asked whether genetic inhibition of MmpS5L5 could restore drug susceptibility of *Rv0678* mutants. To test this, we used CRISPR interference (CRISPRi) to create MmpL5 and MmpS5 CRISPRi knockdowns in both wild-type and *Rv0678* mutant strains (46,47). Transcriptional inhibition of MmpS5 or MmpL5 produced similar effects to the *MmpS5L5* knockout, a >4 fold increased sensitivity to bedaquiline, clofazimine and TBAJ-876 and a smaller increase (0-2 fold) in sensitivity to the DprE1 inhibitor PBTZ-169 (Fig. 2A, compare to Fig. 1B). MmpS5 or MmpL5 CRISPRi knockdown in the *Rv0678* mutant also resulted in increased sensitivity to all substrates, with a smaller 4-fold sensitisation for PBTZ-169 (Fig. 2B). Importantly, for each drug, the MICs were below the corresponding wild-type MICs (Fig. 2B), showing that MmpS5L5 overexpression is the sole cause of drug resistance mediated by *Rv0678* mutants.

**Figure 2.**
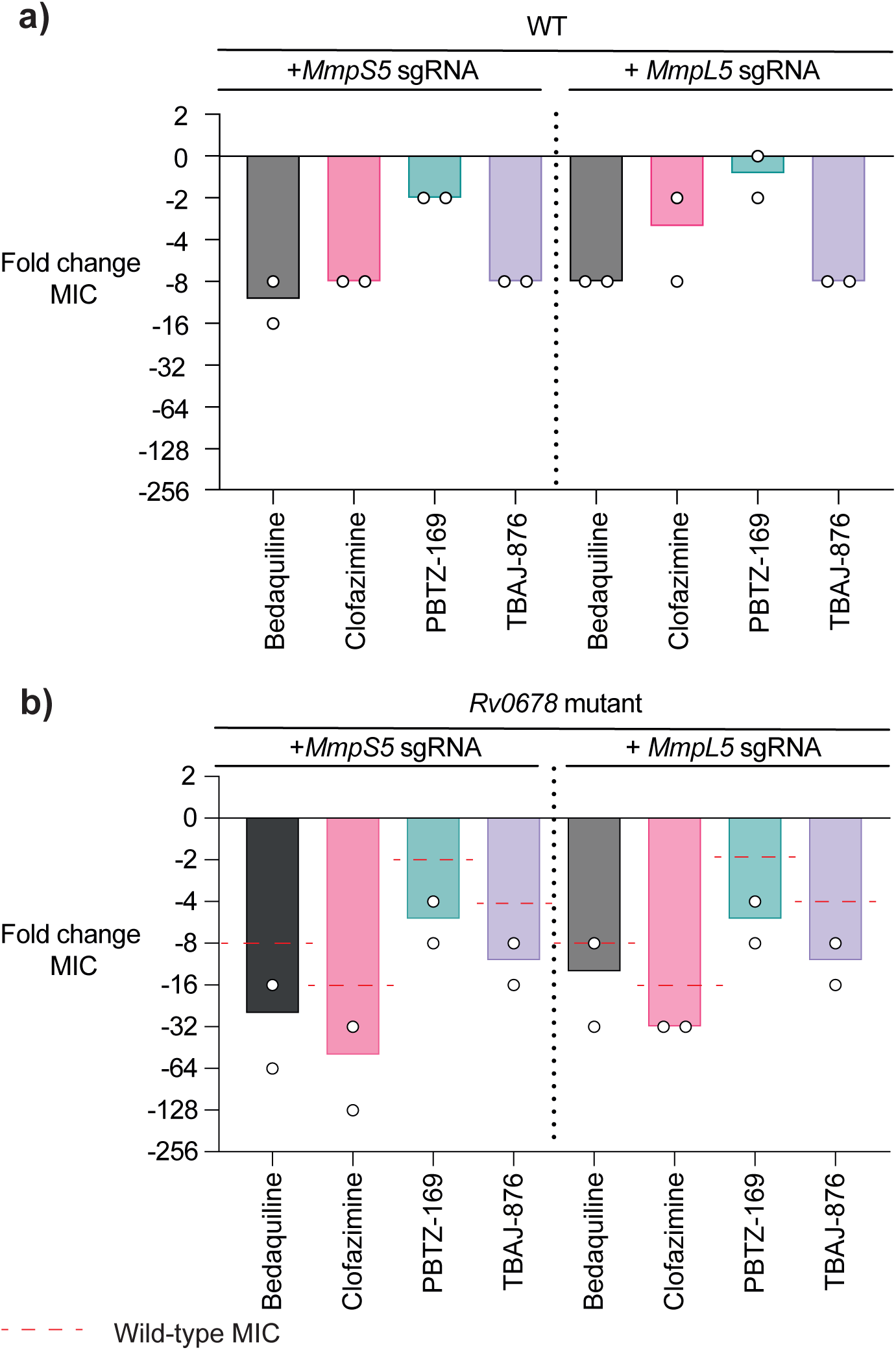
CRISPRi knockdown of MmpS5 or MmpL5 increases drug susceptibility to wild-type and Rv0678 mutant strains of *M. tuberculosis*. (a) Fold change in MICs upon transcriptional knockdown of MmpS5 or MmpL5 in wild-type *M. tuberculosis*. Fold change in MIC is expressed relative to the wild-type parent strain containing a non-targeting CRISPRi plasmid pLJR965. (b) Fold change in MICs upon transcriptional knockdown of MmpS5 or MmpL5 in an *M. tuberculosis Rv0678* background. Fold change in MIC is expressed relative to the *Rv0678* mutant strain containing a non-targeting CRISPRi plasmid pLJR965. Red dotted lines indicate the fold change of the wild-type MIC relative to the *M. tuberculosis Rv0678* mutant. Symbols indicate technical replicates (N = 2). Drug MICs of the WILD-TYPE and Rv0678 mutant are given in *SI Appendix*, Table S2. The differences in MIC fold changes across the tested drugs as compared to those in Figure 1 are due to methodological differences in the MIC assays between the two laboratories performing them.

### Verapamil and its metabolite norverapamil inhibit MmpS5L5 efflux and increase antitubercular drug susceptibility

The finding that genetic knockdown of MmpL5 or MmpS5 increases sensitivity to its substrates even in the context of overexpression, suggested that pharmacological inhibition of MmpS5L5 efflux should also increase activity of these drugs. Verapamil, an inexpensive, orally administered drug used in the treatment of hypertension, cardiac arrhythmias and migraines has been shown to potentiate *M. tuberculosis* activity of multiple drugs, including bedaquiline and clofazimine (48–50). The finding that verapamil increases bedaquiline and clofazimine activity suggests that verapamil inhibits MmpS5L5 pump activity (48,49).

To comprehensively investigate verapamil’s ability to potentiate MmpS5L5-mediated efflux, we performed checkerboard assays with verapamil for multiple MmpL5 substrates (51) (Fig. 3A and *SI Appendix*, Fig. S2). In wild-type bacteria, verapamil showed a dose-dependent decrease in bedaquiline MICs, starting at 6–12.5 µM verapamil (Fig. 3A). Moreover, in the two strains where MmpS5L5 was overexpressed (Rv0678 mutant and complemented MmpS5L5 mutant), verapamil still potentiated bedaquiline sensitivity, starting at 3–6 µM verapamil (Fig. 3A). The wild-type MIC was restored at 12.5–25 µM verapamil. Similar patterns were seen for the bedaquiline derivatives and the other substrates (Fig. 3A). In the MmpS5L5 knockout, verapamil produced only a slight decrease in drug MICs, particularly for bedaquiline, suggesting that it predominantly works by inhibiting MmpS5L5 activity (Fig. 3A). Residual inhibition of the Δ*S5L5* mutant could be due to verapamil’s inhibition of other transporters, as verapamil is able to inhibit multiple classes of transporters (52–55).

**Figure 3.**
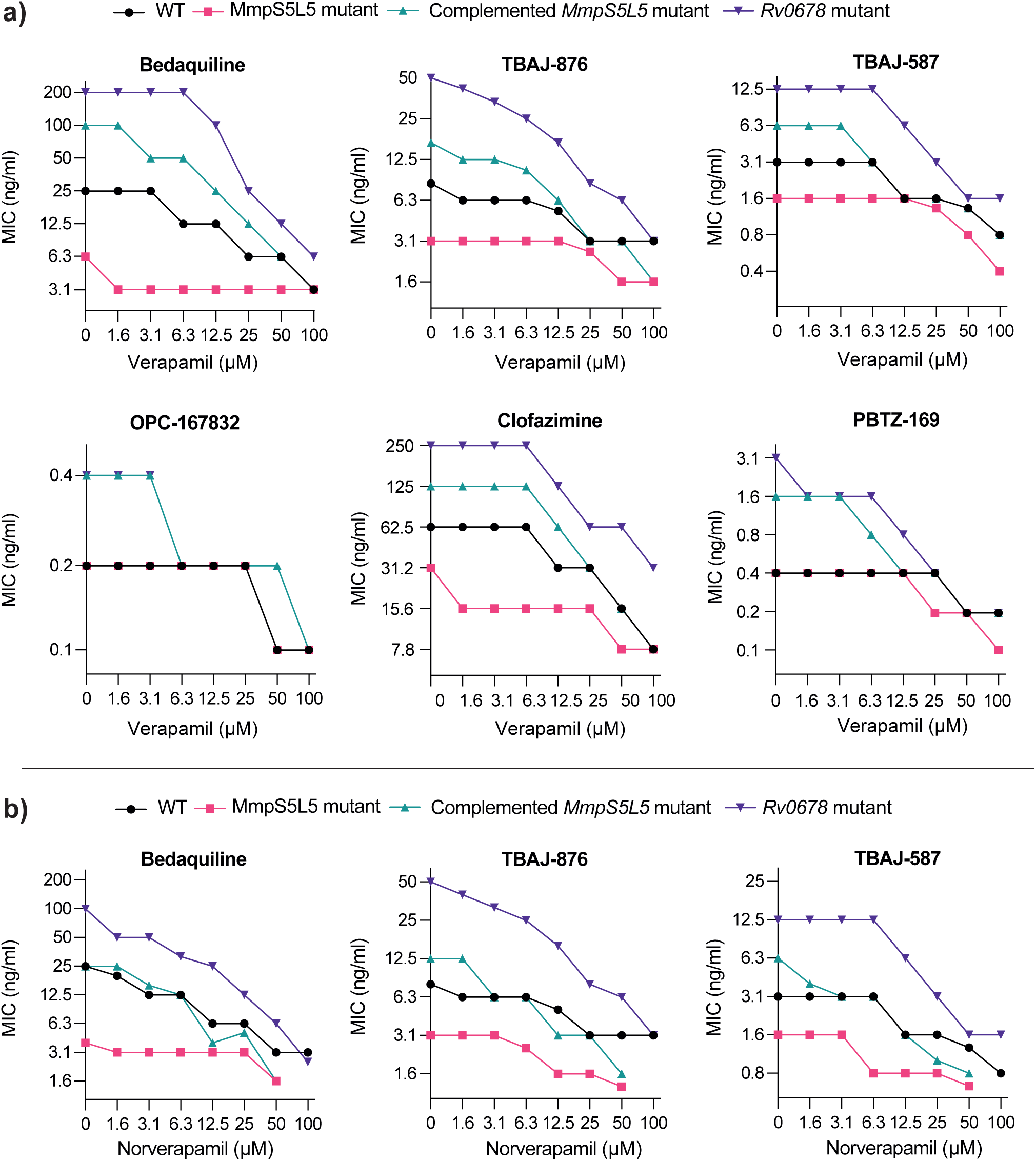
Verapamil and its major metabolite potentiate the activity of MmpL5 substrates in wild-type and MmpS5L5 overexpressing strains (Rv0678 mutant and complemented MmpS5L5 knockout): (a) Isobolograms showing the geometric mean MIC (3 technical replicates) of bedaquiline, clofazimine, PBTZ-169, OPC-167832, TBAJ-876 and TBAJ-587 as a function of verapamil concentration. (b) Isobolograms showing the geometric mean MIC (3 technical replicates) of bedaquiline, TBAJ-876 and TBAJ-587 as a function of norverapamil concentration. Because the norverapamil MIC for *M. tuberculosis* was lower than that of verapamil for the knockout and its complement (100 µM vs. >100 µM), we could not assess its effect beyond 50 µM in these experiments. Note that the Y axis scales differ between graphs.

Whereas verapamil’s benefit in cardiovascular and neurological processes stems from its calcium channel inhibition, it also inhibits human P-glycoprotein (P-gp). Verapamil-caused P-gp inhibition is linked to increased rifampicin susceptibility of *M. tuberculosis* in vitro, and reversal of rifampicin tolerance (56). In vivo, verapamil is rapidly metabolised to norverapamil which has greatly reduced calcium channel inhibition while retaining P-gp inhibition (56–58). Accordingly, we have found that norverapamil inhibits rifampicin efflux with the same potency as verapamil (56). Likewise, in this study, we found that norverapamil potentiated the activity of bedaquiline and its derivatives in a dose-dependent manner with similar potency to verapamil (Fig. 3B).

In previous work, screening a library of approved drugs with incidental P-gp inhibitory activity, we found that the proton pump inhibitor class – omeprazole, lansoprazole, pantoprazole, and rabeprazole - inhibited rifampicin efflux more potently than verapamil, with 2–3 fold greater inhibition at 25 µM and inhibitory activity starting as low as 2.5 µM (56). However, when we tested rabeprazole, we found that it did not potentiate bedaquiline activity at any concentration up to 100 µM of rabeprazole, its MIC for *M. tuberculosis* (*SI Appendix*, Fig. S3). Thus, verapamil is the more promising drug to potentially improve bedaquiline therapy, particularly given that its metabolite norverapamil is also equally effective in vitro.

### Verapamil does not disrupt the protonmotive force used to power MmpSL5 activity

Our findings that verapamil inhibits MmpS5L5 activity extend the body of functional and structural work suggesting that this drug can directly inhibit some efflux pumps (52–56,59–61). However, a prior report using the fluorescent potentiometric dyes 3,3’-dipropylthiadicarbocyanine iodide [DiSC3(5)] and acridine orange (AO) suggested that verapamil exerts this effect indirectly, by dissipating mycobacterial membrane potential (51). MmpL transporters are members of the Resistance-Nodulation-Division (RND) superfamily and are powered by the protonmotive force (PMF) generated during respiration. Therefore, they could be sensitive to molecules such as protonophores and uncouplers that dissipate the PMF. However, there were problems with the data reported and the conclusions drawn in the previous study (51): (i) valinomycin, a potent inhibitor of the membrane potential, was relatively ineffective at collapsing the membrane potential and (ii) when using NADH-energised inverted membrane vesicles (IMVs) of *Mycobacterium bovis* BCG, only very high verapamil concentrations collapsed the ΔpH component of the protonmotive force (PMF) when measured by AO. Specifically, 512 µM verapamil was required to reach values produced by 50 µM of the potent protonophore carbonyl cyanide *m*-chlorophenylhydrazone (CCCP) (51).

We speculated that the use of fluorescent dyes as a qualitative readout of membrane energetics might misrepresent verapamil’s mode of action. To investigate this, we first reproduced the reported AO quenching experiment (51), using IMVs from *M. smegmatis* and measured a significant and concentration-dependent collapse of the transmembrane pH gradient (ΔpH) by verapamil that was equivalent to the positive control CCCP (Fig. 4A). Next, we directly assessed the effect of verapamil on the individual components of the PMF (i.e., the ΔpH and the membrane potential (ΔΨ)), using a quantitative approach with radiometric probes (62). For these experiments, we tested 100 µM verapamil and included the positive controls (i) CCCP that collapses both components, (ii) valinomycin that selectively collapses the membrane potential (ΔΨ), and (iii) nigericin that selectively collapses the transmembrane proton gradient (ΔpH) (62,63). Neither the potassium ionophore valinomycin nor verapamil had any significant effect on the ΔpH in energized cell suspensions compared to the DMSO-treated control when measured using the radioactive probe [7-^14^C] benzoic acid (Fig. 4B). In contrast, both the protonophore CCCP and nigericin collapsed the ΔpH resulting in equilibration of the internal pH with the external pH (pH of medium 6.8) (Fig. 4B). The membrane potential of glucose-energized cell suspensions was in the range 110 – 132 mV and both valinomycin and CCCP collapsed the ΔΨ to values in the range 26 – 60 mV when using [^3^H] methyltriphenylphosphonium iodide (TPP^+^) (Fig. 4C). In contrast, neither nigericin nor verapamil had any significant effect on the membrane potential (Fig. 4C). Another characteristic of protonophoric compounds is their ability to stimulate proton-pumping and activate oxygen consumption by the respiratory chain to uncover spare respiratory capacity (SRC) in an attempt to maintain PMF (64). When the protonophore CCCP was added to glucose-energized cell suspensions of *M. smegmatis*, an immediate activation of the oxygen consumption rate (OCR) was observed with as little as 1 µM CCCP (Fig. 4D). Unlike CCCP, this stimulation of OCR was not observed with increasing concentrations of verapamil (1–10 µM). The same trend was observed in succinate-energized cell suspensions (Fig. 4E) and NADH-energized IMVs (Fig. 4F). Taken together these data demonstrate that verapamil does not disrupt the PMF (membrane potential or ΔpH) or stimulate OCR and therefore does not inactivate the driving force for MmpS5L5 function in mycobacteria.

**Figure 4.**
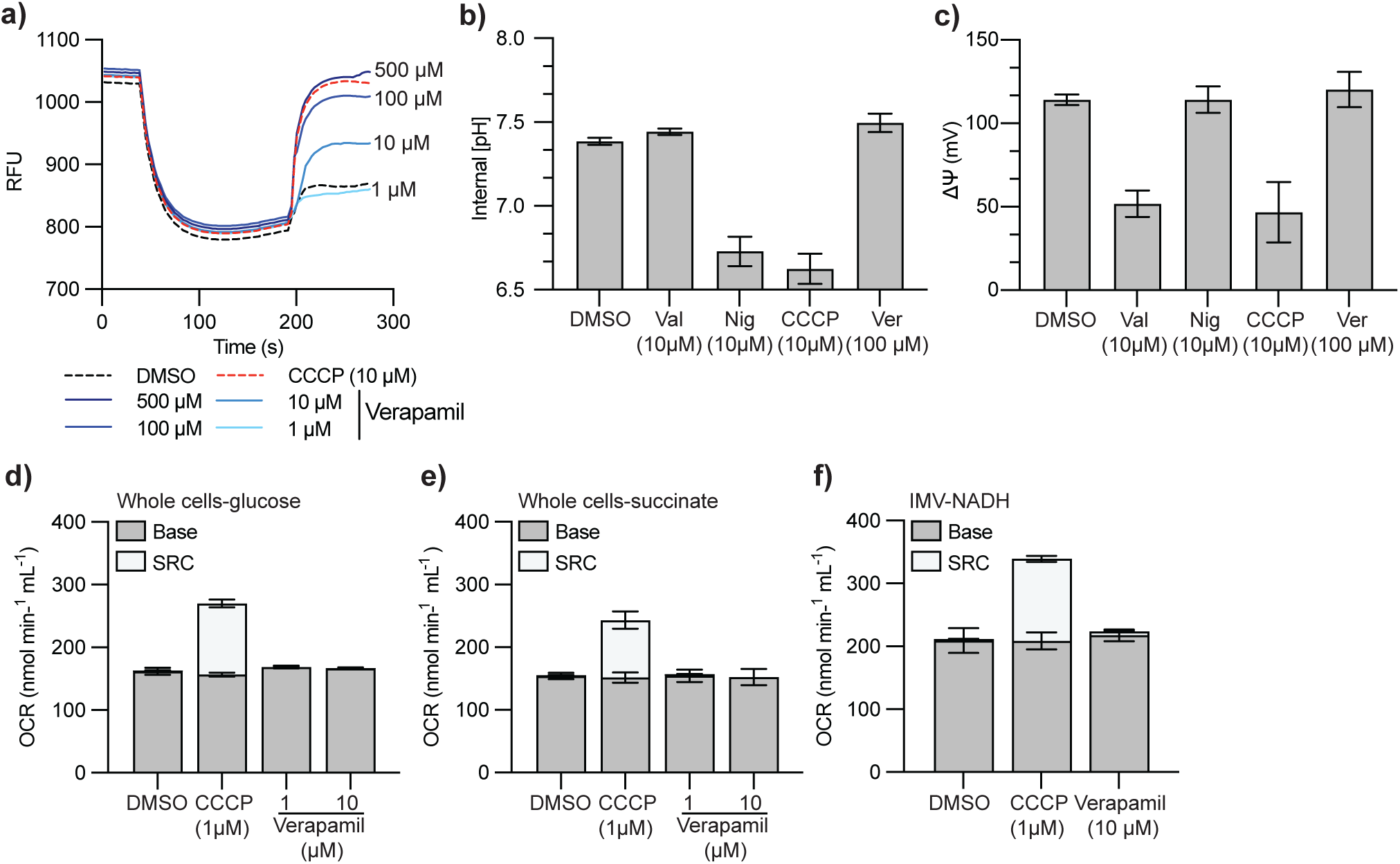
Verapamil does not function as a disruptor of the protonmotive force in mycobacteria: (a) NADH-driven proton translocation in IMVs of M. smegmatis (0.25 mg/mL). Quenching of acridine orange fluorescence in IMVs was initiated with 0.5 mM NADH, and at the indicated time points, either DMSO (control), verapamil (1 – 500 µM) or the protonophore CCCP at 10 µM were added to collapse the proton gradient (reversal of acridine orange fluorescence). IMVs were prepared from M. smegmatis cells grown under aerobic conditions in Hartman’s de Bont [HdB] minimal medium supplemented with 0.2% glucose (w/v). Traces were normalized to a starting value of approximately 1000 relative fluorescence units (RFU) and measurements made with a Cary Eclipse Fluorescence spectrophotometer. The excitation and emission wavelengths were 493 and 530 nm, respectively and experiments are representative of a technical triplicate. (b) Intracellular pH ([7-14C]benzoic acid, 10 µM final concentration) and (c) membrane potential ([3H]methyltriphenylphosphonium iodide, 5 nM final concentration) of glucose-energized cell suspensions (10 mM glucose, OD600 = 1.0) in the presence of either DMSO, valinomycin, nigericin, CCCP or verapamil as indicated. Error bars Indicate standard deviation from three biological replicates. (d-f) The effect of CCCP and verapamil on the oxygen consumption rate (OCR) of either (d) glucose-energized (10 mM final) (e) succinate-energized (10 mM final) cell suspensions of M. smegmatis (OD600 = 1.0) or (e) NADH-energized (0.5 mM final) IMVs of M. smegmatis. OCR was measured using a Oroboros-Oxygraph-2k and inhibitors were added once the OCR reached steady state (∼3 min after electron donor addition). Error bars for OCR measurements represents standard error of the mean (SEM) of three biological replicates and three technical replicates for IMVs.

## Discussion

The emergence and spread of *M. tuberculosis* Rv0678 mutants that upregulate the expression of the MmpS5L5 efflux pump threatens the long-term clinical utility of bedaquiline (65). Here, we have confirmed that the upregulation of MmpS5L5 extends its influence beyond bedaquiline, and provides low-level (2–8 fold) increase in resistance to a spectrum of antitubercular drugs in clinical use or clinical trials. Moreover, we found that MmpS5L5 activity contributes to decreased drug activity even in wild-type strains, suggesting that pharmacological inhibition of this pump could be useful to improve the effectiveness of bedaquiline and other antitubercular drug substrates both in wild-type and Rv0678 mutant strains.

Verapamil, an inexpensive FDA-approved drug that inhibits rifampicin efflux and thereby drug tolerance, has been reported to potentiate bedaquiline and clofazimine activity (7,48,50,51,56,66,67). We have extended these findings to show that verapamil potentiates the activity of all tested MmpS5L5 drug substrates, including the promising, more potent bedaquiline derivatives and DprE1 inhibitors that are currently in clinical trials. Previous work suggested that verapamil might inhibit drug efflux by acting as an uncoupler of the PMF (51). The chemical nature of verapamil makes it highly unlikely that it could function as a disruptor of the PMF i.e., exhibit protonophore-like activity, as its positive charge cannot be delocalized, making it energetically unfavourable for membrane entry. All charged protonophores have highly delocalized charges (68,69). Verapamil has also been shown to directly bind and inhibit multiple efflux transporter families (53–55,59), including the ABC-transporter P-gp whose activity does not require the PMF, suggesting that verapamil acts directly and not by disrupting the PMF. In this work, we conclusively rule out that verapamil’s inhibition of efflux is due to it functioning as a disruptor of the PMF in mycobacteria (51). Rather, our results point to verapamil directly inhibiting MmpS5L5 activity as it has only limited bedaquiline potentiation in the MmpS5L5 pump knockout.

As is the case for inhibition of rifampicin efflux (50,56), we find that verapamil’s major metabolite norverapamil potentiates the efficacy of bedaquiline and its derivatives. This finding is important because norverapamil’s plasma levels are higher than verapamil’s (70–74). Therefore, it could substantially augment verapamil’s effects on inhibiting MmpS5L5 activity. Verapamil is supplied as a racemic mixture and the predominant R-enantiomer has substantially less cardiac activity but similar P-gp and rifampicin efflux inhibition to the S-enantiomer (56). Therefore, both enantiomers are likely to contribute to MmpS5L5 efflux inhibition. Animal studies have shown that verapamil and norverapamil concentrate in lung tissue at levels 40-fold higher than that found in blood (75,76), showing that potentiating concentrations of verapamil and norverapamil may be achievable in the human lung (77), the primary site of TB. Verapamil may have an effect on bedaquiline pharmacokinetics through its inhibition of CYP3A4 (78–80), which is the primary cytochrome P450 enzyme that converts bedaquiline to the less active M2 metabolite (81). Verapamil, including its enantiomers and metabolites, all inhibit *M. tuberculosis* growth in macrophages even in the absence of antibiotics, adding further to its therapeutic potential (56,66). Finally, verapamil potentiates bedaquiline in murine infection models resulting in a greater reduction in bacterial burden (7,48,67). Together, these properties and effects of verapamil improve its prospects of boosting the clinical effectiveness of bedaquiline.

The rapid emergence of bedaquiline resistance warrants urgent therapies to prevent and treat this. Verapamil is an ideal candidate given its availability as an inexpensive generic medicine, decades-long worldwide clinical use, safety, well-characterized pharmacology, and adverse effect profile (82). This work suggests that verapamil could be tried as adjunctive therapy to both prevent and treat resistance to bedaquiline and other MmpS5L5 substrates. At the same time, we hope that it will spur the urgent development of potent MmpS5L5 inhibitors for co-administration in bedaquiline-containing regimens.

## Methods

### Bacterial strains and growth conditions

The *M. tuberculosis* strain mc^2^6206 (H37Rv Δ*panCD*, Δ*leuCD*) is an auxotrophic derivative of H37Rv (83) and was obtained from W.R. Jacobs, Jr. *M. tuberculosis* and resistant derivatives were grown at 37°C in Middlebrook 7H9 liquid medium or on 7H10 solid medium supplemented with OADC (0.005% oleic acid [Sigma, #O1008], 0.5% BSA [pH Scientific, #PH100], 0.2% dextrose [Sigma, #G8270], 0.085% catalase [Sigma, #02071]), 25 µg/ml pantothenic acid (Sigma, #21210), 50 µg/ml L-leucine (Sigma, #L8000) and when required 20 µg/ml kanamycin. Liquid cultures were maintained in T25 culture flasks (Sigma, #156367) and supplemented with 0.05% tyloxapol (Sigma, #T8761). 7H9 refers to fully supplemented 7H9 media, whilst 7H9-K refers to 7H9 supplemented media with kanamycin. *Escherichia coli* strain MC1061 was used for the cloning of CRISPRi plasmids. *E. coli* MC1061 was grown at 37°C in LB or on LB-agar supplemented with kanamycin at 50 µg/ml. Compounds used in susceptibility assays are as follows: bedaquiline (MedChemExpress #HY-14881), clofazimine (Sigma, #28266), PBTZ-169 (Cayman Chemical, #22202), TBAJ-876 (Gift courtesy of TB-Alliance), TBAJ-587 (MedChemExpress #HY-111747), OPC-167832 (MedChemExpress #HY-134940). All compounds were dissolved in DMSO.

### Construction and complementation of MmpS5L5 deletion mutant

MmpS5L5 knockout strains in *M. tuberculosis* were generated by recombineering (84). *M. tuberculosis* was transformed with pNIT-ET-SacB-Kan. pNIT-ET-SacB-Kan was a gift from Kenan Murphy (Addgene plasmid #107692). Single colonies of the pNIT containing strain were inoculated into 50 ml 7H9 media supplemented with 25 μg/ml kanamycin and grown at 37 °C until an OD600 of ∼0.8 was reached. Expression of Che9c RecET enzymes was induced by addition of isovaleronitrile to 1 μM final concentration and incubated for 8 hours at 37 °C before addition of sterile-filtered 2 M glycine to a final concentration of 0.2 M. The induced cell suspensions were further incubated for 16 hours at 37°C before preparing electrocompetent cells.

BsaI-domesticated pKM342 (adapted from Addgene #71486) containing 500 bp homology arms amplified from flanking regions of the *M. tuberculosis* MmpS5L5 operon was digested for 16 hours with EcoRV (NEB #R1095). Digested plasmid DNA was purified with a PCR clean-up kit. An amount of DNA corresponding to 1 μg hyg cassette fragment was electroporated into induced, freshly prepared electrocompetent cells and plated on 7H10 agar supplemented with 10% (v/v) OADS, 50 μg/ml L-leucine, 24 μg/ml D-pantothenic acid and 50 µg/ml hygromycin. Plates were incubated at 37 °C for 28 days. Hygromycin-resistant colonies were inoculated into 7H9 media supplemented with 50 μg/ml hygromycin and once growth reached mid-log phase, crude genomic DNA was isolated. Isolates were screened for correctly inserted DNA constructs by PCR using primers designed to span the junction between the integrated knockout construct and flanking genome regions.

To generate an unmarked mutant strain, pCre-SacB-Zeo (Addgene #107706) was transformed into *M. tuberculosis* Δ*S5L5*::*hyg* and plated on 7H10 agar supplemented with 10% (v/v) OADS, 50 μg/ml L-leucine and 24 μg/ml D-pantothenic acid and 25 μg/ml zeocin (Invitrogen, #R25001). The pNIT-SacB-RecET plasmid is incompatible with pCre-SacB-Zeo, so curing pNIT-SacB-RecET before this step was unnecessary. Zeocin-resistant colonies were inoculated into 7H9 media supplemented with 25 μg/ml zeocin and incubated at 37 °C. Isolates were screened for excision of the hyg cassette by PCR across the *loxP-hyg-loxP* sequence and resolving the PCR products by agarose gel electrophoresis. Sensitivity to hygromycin was confirmed by streaking each excised isolate on 7H10 agar supplemented either with 50 μg/ml hygromycin or 10% (w/v) sucrose (to cure the pCre-SacB-Zeo plasmid). Genomic DNA was isolated from hygromycin-sensitive isolates and Sanger sequencing of a PCR product spanning the locus was done to confirm correct deletion and excision of the *MmpS5L5* operon.

### Construction and transformation of CRISPRi plasmids that target genes of interest

sgRNA sequences that target *mmpS5* or *mmpL5* were cloned as previously described (47). Briefly, the sgRNA sequence and a complementary sequence were ordered with GGGA and AAAC overhangs. Oligos were annealed and cloned into pLJR965 using BsmBI and confirmed using Sanger sequencing as previously described. All ordered oligos and constructed CRISPRi plasmids are listed in Table 2. CRISPRi plasmids were transformed into *M. tuberculosis* strains as previously described.

### Minimum inhibitory concentration (MIC) determination (Laboratory 1)

MIC assays were conducted using the broth microdilution method. MIC plates were prepared with 2X concentration of test compound in Middlebrook 7H9 + 10% OADS without tween-80 in polystyrene round-bottom 96-well plates (Corning, Falcon #351177). Compounds were dissolved in anhydrous DMSO and stored in single-use aliquots at −70 °C. For plate preparation, a 100X stock of the highest antimicrobial concentration to be tested was prepared in DMSO, and from this a ten-point two-fold dilution series was made using DMSO as the diluent. Each point in the dilution series was further diluted 50-fold in 7H9 + OADS without Tween 80 to produce a final 2X concentration of compound in media and 100 µl per well was dispensed column-wise. Media only and DMSO control columns were included. Plates were sealed with film and stored at −20°C. After thawing and prior to use, plates were centrifuged at 500 x g for 5 minutes. Test strains were diluted to 0.5 McFarland (McF) units in 7H9 + 10% OADS with 0.05% Tween 80 using a nephelometer. 100 µl 0.5 McF cell suspension was diluted 100-fold in 7H9 + OADS + 48 µg/ml pantothenic acid + 100 µg/ml L-leucine + 0.05% Tween 80. 100 µl of this diluted bacterial suspension, corresponding to ∼ 5 x10^5^ CFU, was added to each well of the 96-well plate to obtain a 1X concentration of test compound. The concentration of DMSO after addition of bacteria was 1%. Plates were sealed with tape and incubated at 37°C for 14 days. Plates were imaged using an Epson perfection V850 photo scanner. Minimum inhibitory concentrations were defined as the minimum concentration of test compound that results in no growth visible by eye in scanned images.

### Minimum inhibitory concentration assays (Laboratory 2)

The susceptibility of the drug-susceptible-parent or resistant strains to different compounds was determined using Minimum inhibitory concentration (MIC) assays as previously described (19,85). Briefly, 75 µl 7H9 media was added to the inner wells (rows B–G, columns 3–11) of a 96-well flat-bottomed microtiter plate (ThermoFisher Scientific), with 150LJµl 7H9 media being added to outer wells and left as media only controls. 150 µl of 7H9 media containing compound of interest at the required starting concentration was added to column 2 of row B–G and diluted 2-fold down to column 10. Column 11 was kept as solvent only. 75 µl of culture diluted to an OD_600_ of 0.01 was added to inner wells of the 96-well microtiter plate to achieve a starting OD_600_ of 0.005. Plates were incubated for 10 days at 37°C without shaking. After 10 days, plates were covered with plate seals, shaken for 1LJmin and the OD_600_ was determined using a Varioskan Flash microplate reader (ThermoFisher Scientific). OD_600_ reads from duplicate plates were corrected for the background OD_600_ and expressed as values relative to the growth of the no-compound control. The MIC was defined here as the IC_90_, the concentration of drug that results in >90% growth inhibition, which aligns with the visual MIC determination method used by laboratory 1.

### Compound susceptibility and viability assays against *M. tuberculosis* strains pre-depleted for genetic targets (Laboratory 2)

*M. tuberculosis* strains depleted for genes of interest using CRISPRi were prepared as previously described (86). Briefly, log phase culture was diluted to an OD_600_ of 0.005 in 10 ml of 7H9-K with 300 ng/ml ATc in T25 tissue culture flasks standing upright and grown without shaking at 37°C for 5-days to pre-deplete target genes. After 5-days culture was diluted to a OD_600_ of 0.01. Assay plates for susceptibility assays were prepared as described above in 7H9-K+ATc. 75 µl of OD_600_ adjusted culture was added to assay plates. Plates were incubated at 37°C for 10 days without shaking. After 10 days, plates were covered with plate seals, shaken for 1 min and the OD_600_ was determined using a Varioskan Flash microplate reader (ThermoFisher Scientific). OD_600_ readings from duplicate plates were corrected for background, and values relative to the growth of the no compound control were analysed. The MIC was defined here as the IC_90_: the concentration of drug that results in >90% growth inhibition.

### Immunoblotting

50 ml OD_600_ = 1.0 cell suspensions were harvested by centrifugation at 4,000 x g and washed twice with PBS + 0.1% Tween 80. The cell pellet was resuspended in 1.0 ml lysis buffer (50 mM HEPES pH 8.0, 150 mM NaCl, 10% glycerol) and transferred to 2 ml lysing matrix B tubes (MP Biomedical). Cell lysis was performed using beat-beating using a FastPrep homogeniser (MP Biomedical) for 6 cycles of 45 seconds at 6 m/s, with a wait time of 3 minutes. Unbroken cells and debris were pelleted at 1,000 x g. Protein concentration was measured using a bicinchoninic acid (BCA) assay (Thermo-Scientific) kit according to the manufacturer’s instructions, using bovine serum albumin standards (Thermo-Scientific #23208) as a calibration curve. Proteins were resolved by SDS-PAGE before being transferred to 0.2 µm PVDF membranes (BioRad #1704156) by semi-dry blotting using a Transblot turbo transfer system (BioRad). Membranes were blocked using 3% (w/v) skimmed milk powder (Sigma #70166) in PBS. Membranes were incubated with primary antibodies at appropriate dilutions in 3% (w/v) skimmed milk powder in PBS for 1 hour at 22 °C or overnight at 4 °C. Membranes were washed using PBS supplemented with 0.1 % (v/v) Tween 20. Membranes were incubated with secondary antibodies at appropriate dilutions in 3% (w/v) skimmed milk powder in PBS for 1 hour at 22°C or overnight at 4°C and washed with PBS supplemented with 0.1% (v/v) Tween 20. Blots were imaged at 700 and 800 nm using a LI-COR Odyssey imaging system. 3xFLAG tags on MmpL5 were detected using a monoclonal mouse anti-FLAG M2 antibody (Sigma-Aldrich #F1804) at 1 µg/ml (1:1000). Goat anti-mouse IgG conjugated with Dylight™ 800 4x PEG (Cell signalling technology #5257) was used at 0.1 µg/ml (1:10,000). MmpS5 was detected with a custom rabbit polyclonal antibody (Genscript) at 0.1 µg/ml (1:5,000). Goat anti-rabbit IgG conjugated with Dylight™ 800 4x PEG (Cell signalling technology #5151) was used at 0.1 µg/ml (1:10,000).

### Checkerboard assay

Checkerboard assays were performed by preparing 2x final concentrations of Verapamil or Norverapamil in 7H9 + OADS + 24 µg/ml pantothenate, 50 µg/ml L-leucine + 0.05% Tween-80. 0.5 McF bacterial suspensions, prepared as previously, were added at a ratio of 1:100 to each tube containing media with 2x final concentration of verapamil/norverapamil. 100 µl of each dilution containing ∼ 1x 10^5^ CFU bacteria was added row by row, into the MIC plates for each drug. 7 x 2-fold dilutions starting from 100 µM were used. One row was a no verapamil / norverapamil control.

### Measurement of the protonmotive force and oxygen consumption rate in mycobacteria

*M. smegmatis* strain mc^2^155 (87) was routinely grown at 37°C with agitation (200 rpm) in Hartman’s de Bont (HdB) minimal medium (88) supplemented with 0.05% Tween 80 (Sigma-Aldrich) and 0.2% glucose. Preparation of inverted membrane vesicles (IMVs) and washed cell suspensions were carried out as previously described (89). The membrane potential and transmembrane pH gradient determinations and proton-pumping assays were performed as previously described (89). The oxygen consumption rate (OCR) of either energized cell suspensions or IMVs was measured using an Oroboros-Oxygraph-2k instrument and inhibitors were added once the OCR reached a steady state (∼3 min after electron donor addition).

## Acknowledgments and Funding Sources

We thank all Cook and Ramakrishnan lab members and Departmental colleagues for helpful discussions. This research was financially supported by the Maurice Wilkins Centre for Molecular Biodiscovery and the Marsden Fund (Royal Society of New Zealand) (grant number UOO1807)(G.C,) and a Wellcome Trust Principal Research Fellowship (223103/Z/21/Z) and NIH MERIT award R37 AI054503 (L.R.). NW was supported by a University of Otago Doctoral Scholarship. MM was supported by the Health Research Council of New Zealand Sir Charles Hercus Health Research Fellowship (22/156).

