## Supplementary Information for "Verapamil and its metabolite norverapamil inhibit the *Mycobacterium tuberculosis* MmpS5L5 efflux pump to potentiate the activity bedaquiline and other antitubercular drugs"

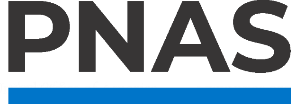

**Supporting Information for**

**Verapamil and its metabolite norverapamil inhibit the *Mycobacterium tuberculosis* MmpS5L5 efflux pump to potentiate the activity bedaquiline and other antitubercular drugs**

Adam J Fountain, Natalie JE Waller, Chen-Yi Cheung, William Jowsey, Michael T Chrisp, Mark Troll, Paul H Edelstein, Gregory M Cook*, Matthew B McNeil* and Lalita Ramakrishnan*

Gregory Cook

Matthew McNeil

Lalita Ramakrishnan

**This PDF file includes:**

Figures S1 to S3

Tables S1 to S2

SI References

Supporting Information

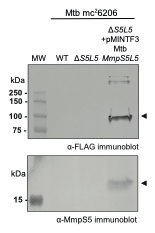

**Fig S1. Immunoblot analysis of complemented strains:**

20 µg whole cell lysate from Mtb mc^2^6206, ∆*S5L5*::loxP and ∆*S5L5*::loxP attB_L5_::pMINTF3 Mtb MmpS5L5 strains probed with mouse anti-FLAG or anti-Mtb MmpS5 (VFADDPEPFDPKVVC) antibody. Endogenous levels of MmpS5 in the wild-type strain are undetectable in whole-cell lysate.

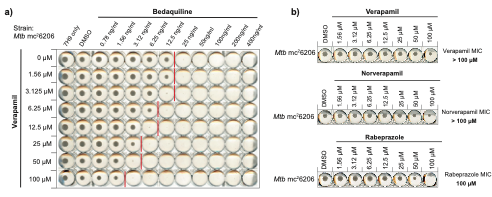

**Fig S2. Representative checkerboard**

(a) Representative checkerboard of BDQ–VER in Mtb mc^2^6206. Red lines indicate drug MICs. (b) Verapamil, norverapamil and rabeprazole only columns, showing that verapamil/norverapamil MIC > 100 µM, whilst rabeprazole MIC = 100 µM.

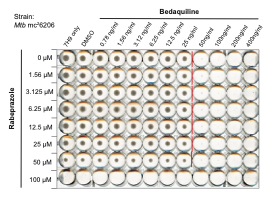

**Fig S3. Rabeprazole does not potentiate bedaquiline MIC**

(a) Representative checkerboard of BDQ–Rabeprazole in Mtb mc^2^6206. Red lines indicate drug MIC. Rabeprazole has an MIC of 100 µM.

**Table S1. Strains, plasmids and primers used in this study.**

*E.coli strains*

| Strain | Description | Source |
| --- | --- | --- |
| MC1061 | Cloning strain for CRISPRi plasmids | Lab strain |
| DH5-α | Cloning strain for knockout and complementation plasmids | Invitrogen |

*M. tuberculosis* and *M. smegmatis* strains

| Strain | Description | Source |
| --- | --- | --- |
| *M. tuberculosis* mc^2^6206 | Drug-susceptible | (1) |
| *M. tuberculosis* mc^2^6206 *Rv0678*^G65GfsX10^ | Isolated from bedaquiline containing media | (2) |
| *M. tuberculosis* mc^2^6206 ∆*S5L5*::loxP | Unmarked MmpS5L5 KO strain | This study |
| *M. tuberculosis* mc^2^6206 ∆*S5L5*::loxP, attB_L5_::pMINTF3 Mtb MmpS5L5 | Unmarked MmpS5L5 KO strain complemented with pMINTF3 Mtb MmpS5L5, Kan^R^. | This study |
| *M. smegmatis* mc^2^155 | Transformable lab strain of *M.smegmatis* | (3) |

Plasmids

| Plasmid name | Description | Target sequence (Coding, 5’-3’) | PAM (non coding 5’-3’, NN…...) | Source | |
| --- | --- | --- | --- | --- | --- |
| pJLR965 | Cloning plasmid and negative control |  |  | Addgene #115163 | |
| pCi1214 | CRISPRi knockdown of MmpL5 | CTGAGCTTCACCCGACTGCC | ACAGAAC | This study | |
| pCi1215 | CRISPRi knockdown of MmpS5 | GGTTCCGAAGGCATCTTGGT | AAAGAAA | This study | |
| pNIT-ET-SacB-Kan | Episomal Isovaleronitrile-inducible expression of phage Che9c RecET enzymes for recombineering | | | | Addgene #107692 |
| pCre-SacB-Zeo | Episomal expression of Cre recombinase for excision of loxP-hyg-loxP casette | | | | Addgene #107706 |
| pKM342-S5L5KO | BsaI-domesticated pKM342 (Adapted from Addgene #71486) containing ~500bp sequences upstream and downstream of MmpS5L5 operon flanking a loxP-hyg-loxP cassette for recombineering. | | | | This study |
| pMINTF3 Mtb MmpS5L5 | L5 int, L5 attP integrating vector, constitutively expressing MmpS5L5-3xFLAG from pmyctetO promoter | | | This study | |

Oligonucleotides used in this study

| Oligos for CRISPRi plasmid construction | |
| --- | --- |
| Oligo Name | Oligo Sequence |
| mmpL5_a_TB_Fcs | GGGAGGCAGTCGGGTGAAGCTCAG |
| mmpL5_a_TB_Rcs | AAACCTGAGCTTCACCCGACTGCC |
| mmpS5_a_TB_Fcs | GGGAACCAAGATGCCTTCGGAACC |
| mmpS5_a_TB_Rcs | AAACGGTTCCGAAGGCATCTTGGT |

**Table S2. Laboratory 1 and 2 MIC values**

Laboratory 1 MIC values (Figures 1 and 3)

|  | Bedaquiline | | Clofazimine | | PBTZ-169 | | OPC-167832 | | | TBAJ-876 | | | TBAJ-587 | | |
| --- | --- | --- | --- | --- | --- | --- | --- | --- | --- | --- | --- | --- | --- | --- | --- |
| Strain | Geometric mean MIC (ng/ml) | Range (ng/ml) | Geometric mean MIC (ng/ml) | Range (ng/ml) | Geometric mean MIC (ng/ml) | Range  (ng/ml) | Geometric mean MIC (ng/ml) | Range  (ng/ml) | Geometric mean MIC (ng/ml) | | Range  (ng/ml) | Geometric mean MIC  (ng/ml) | | | Range (ng/ml) |
| Mtb H37Rv mc^2^6206 | 25 | 25 | 198 | 125–500 | 0.5 | 0.39–0.79 | 0.195 | 0.195–0.39 | 7.9 | | 6.25–12.5 | 3.1 | | 3.1 | |
| Mtb H37Rv mc^2^6206 Rv0678(G65GfsX10) | 141 | 100–200 | 500 | 500 | 3.12 | 3.12 | 0.491 | 0.39–0.78 | 44.5 | | 25–50 | 12.5 | | 12.5 | |
| Mtb H37Rv mc^2^6206 ∆*S5L5*::loxP | 6.25 | 6.25 | 31.2 | 31.2 | 0.39 | 0.39 | 0.195 | 0.195 | 3.12 | | 3.12 | 1.56 | | 1.56 | |
| Mtb H37Rv mc^2^6206 ∆*S5L5*::loxP  attB_L5_::pMINTF3 *MmpS5L5* | 84 | 50–100 | 250 | 250 | 1.56 | 1.56 | 0.491 | 0.39–0.78 | 14 | | 12.5–25.0 | 6.25 | | 6.25 | |

Laboratory 2 MIC values (Figure 2)

|  | Bedaquiline | | | Clofazimine | | | PBTZ-169 | | | TBAJ-876 | | |
| --- | --- | --- | --- | --- | --- | --- | --- | --- | --- | --- | --- | --- |
| Strain | | Geometric mean MIC (ng/ml) | Range  (ng/ml) | | Geometric mean MIC (ng/ml) | Range  (ng/ml) | | Geometric mean MIC (ng/ml) | Range  (ng/ml) | | Geometric mean MIC (ng/ml) | Range  (ng/ml) |
| Mtb H37Rv mc^2^6206 | | 148 | 88-216 | | 135.9 | 11.5–370 | | 0.25 | 0.18–0.71 | | 36.3 | 25.6–51.3 |
| Mtb H37Rv mc^2^6206 + MmpS5 gRNA | | 27 | 27 | | 23.1 | 23.1 | | 1.3* | 0.9–1.78 | | 6.4 | 6.4 |
| Mtb H37Rv mc^2^6206 + MmpL5 gRNA | | 19 | 13.5–27 | | 16.4 | 11.5–23.1 | | 0.9* | 0.9 | | 6.4 | 6.4 |
| Mtb H37Rv mc^2^6206 Rv0678(G65GfsX10) | | 796 | 436–1736 | | 737.0 | 182–1439 | | 1.6 | 0.7–2.9 | | 290 | 102–411 |
| Mtb H37Rv mc^2^6206 Rv0678(G65GfsX10) + MmpS5 gRNA | | 75 | 54–108 | | 5.8 | 5.8 | | 0.18 | 0.18 | | 9.1 | 6.4–12.8 |
| Mtb H37Rv mc^2^6206 Rv0678(G65GfsX10) + MmpL5 gRNA | | 38 | 27–54 | | 2.9 | 1.4–5.8 | | 0.18 | 0.18 | | 9.1 | 6.4–12.8 |

Note: Differences in MICs between laboratories 1 and 2 are likely a result of differing methods of MIC determination. For clarity, MICs values are expressed in ng/ml rather than the standard unit of µg/ml. * - This experiment had higher MICs across all strains
